# Lung injury induces alveolar type 2 cell hypertrophy and polyploidy with implications for repair and regeneration

**DOI:** 10.1101/2021.08.10.455833

**Authors:** Anthea Weng, Mariana Maciel-Herrerias, Satoshi Watanabe, Lynn Welch, Annette S. Flozak, Rogan Grant, Raul Piseaux Aillon, Laura Dada, Seung Hye Han, Monique Hinchcliff, Alexander Misharin, GR Scott Budinger, Cara J. Gottardi

## Abstract

Epithelial polyploidization post-injury is a conserved phenomenon, recently shown to improve barrier restoration during wound healing. Whether lung injury can induce alveolar epithelial polyploidy is not known. We show that bleomycin injury induces AT2 cell hypertrophy and polyploidy. AT2 polyploidization is also seen in short term ex vivo cultures, where AT2-to-AT1 trans-differentiation is associated with substantial binucleation due to failed cytokinesis. Both hypertrophic and polyploid features of AT2 cells can be attenuated by inhibiting the integrated stress response (ISR) using the small molecule ISRIB. These data suggest that AT2 hypertrophic growth and polyploidization may be a feature of alveolar epithelial injury. As AT2 cells serve as facultative progenitors for the distal lung epithelium, a propensity for injury-induced binucleation has implications for AT2 self-renewal and regenerative potential upon reinjury, which may benefit from targeting the ISR.

## Introduction

Genome wide association and classical genetic studies of families with pulmonary fibrosis implicate disordered epithelial repair as a driver of lung fibrosis (1–3). While this hypothesis has been confirmed in mouse models of fibrosis that localize fibrosis-promoting effects of global mutations to the lung epithelium (4–6), the sequence of cellular events that drive fibrosis persistence remains unclear. Understanding how the distal lung epithelium repairs after injury is also relevant to patients with severe lung injury induced by SARS-CoV-2 pneumonia, where some will develop persistent and progressive lung fibrosis (7–12). Thus, while it is clear surfactant producing-alveolar type 2 pneumocytes (AT2 cells) are progenitors that give rise to alveolar type 1 pneumocytes (AT1 cells)(13, 14), the variability between individuals in the AT2-to-AT1 cell trans-differentiation program to restore normal alveolar function is unexplained.

Pathologists have long recognized that a range of injurious stimuli lead to the accumulation of abnormally large AT2 cells, described as “hypertrophic or hyperplastic” AT2 cells (15–18). Morphologically similar cells are seen in autopsy specimens or lung explants from patients with lung injury secondary to SARS-CoV-2 pneumonia (7, 11, 19), but mechanisms underlying this cell state and its consequences for lung repair post-injury remain puzzling. A recent convergence of single cell transcriptomic-studies from a range of mouse injury models suggests that instead of fully progressing to an AT1 cell fate, AT2 cells become stalled at an intermediate transition-state characterized by activation of cell cycle arrest pathways (p16, p21, p53),TGFβ signaling and matrix remodeling (SERPINE1)(20–23). Serially collected single cell RNA-sequencing data over the course of bleomycin induced injury shows the formation of this intermediate transition state is preceded by increased expression of genes in the integrated stress response (ISR)(22). In tissue sections, these transitional cells are characterized by elevated levels of specific intermediate filament keratins (KRT8/18 in mice; KRT17 in humans) and other structural proteins, along with an intermediate AT2-to-AT1 morphology (22, 23). These data suggest small cuboidal AT2 cells differentiate into large and flattened AT1 cells through a morphologically stressful transition that appears prone to stalling after injury. Importantly, recent evidence suggests the AT2-to-AT1 transition state is not fixed, as therapeutic targeting of p53 or ISR signaling appears to guide AT2 cells through this vulnerable trans-differentiation step (23, 24). However, the cellular processes most vulnerable to this morphogenetic transition, which drive persistence of the AT2-to-AT1 transition state in injured lungs and can be reversed by these therapies are unknown.

Epithelial polyploidization and consequent hypertrophy post-injury is a conserved phenomenon observed from flies to vertebrates (25). It is a process by which cells increase their DNA content and bypass mitosis. Polyploid cells are larger than their diploid counterparts because cell size scales with DNA content (26). Cell hypertrophy is advantageous during epithelial repair, as larger cells manifest less junctional surface area per unit of epithelium, leading to decreased permeability in an otherwise leaky-barrier environment (27, 28). Increased DNA copies can also favor adaptation to cell and environmental stress, in ways that are context and injury dependent (29–31). However, these benefits of polyploidization come at a cost, as polyploid cells will less faithfully segregate DNA during mitosis (32–34). Thus, polyploidization of differentiated cell types—particularly those that serve also as progenitors (e.g., AT2 cells)—is likely to adversely impact future regenerative potential of the tissue. Whether AT2 polyploidy underlies injury-induced AT2 hypertrophy, or is a consequence of hypertrophic growth, and drives persistence of the AT2-to-AT1 cell stalled state is unknown.

In this study, we show that lung injury leads to AT2 hypertrophy and polyploidization—notably binucleated AT2s—during the acute injury response. *Ex vivo* analysis suggests the route to polyploidy is via failed cytokinesis during the AT2-to-AT1 flattening process. A small molecule inhibitor of the integrated stress response, ISRIB, which we have previously shown ameliorates fibrosis development in murine models (24), inhibits the abundance of hypertrophic and polyploid AT2 cells. These data suggest that modulating stress signals during the morphogenetically challenging AT2-to-AT1 transition may improve key steps through which injured AT2 cells divide and differentiate to restore the alveolar epithelium. Implications of failing to remove and replace polyploid AT2 cells after lung injury are discussed.

## Results

### Lung injury induces AT2 hypertrophy

To understand how lung injury alters the morphogenetic sequence by which AT2 cells grow, divide and differentiate to an AT1 fate, we used a genetic lineage tracing system to permanently label AT2 cells in *Sftpc*^Cre-ER^; ^Lox-Stop-Lox^zsGreen mice subjected to the bleomycin model of injury and fibrogenic repair (24) (Fig.1A). As with previous studies (15–18), we confirm that bleomycin injury leads to zsGreen+ cells that are larger than AT2 lineage-labeled mice exposed to saline (Fig. 1B and C, quantification from 2 mice, 5 fields of view; Frozen section image analysis pipeline in Fig. S1A). Cell size difference may be more readily apparent in 14μm thick frozen sections than thinner 4μm paraffin sections, likely because the former method favors cell visualization along a greater portion of the curved alveolus (Fig. S1B). The injury-induced size increase is not an aberrant feature of cells expressing the fluorescence lineage-label, as AT2 ceIls lacking zsGreen but expressing surfactant protein C (SpC) also increase in size (Fig. S1C). Since standard sectioning (i.e., planimetry) is known to underestimate AT2 cell expansion post-injury (35, 36), we also confirmed AT2 hypertrophic growth using cleared, thick-section (200μm) confocal imaging with 3-dimensional reconstruction (Fig. 1D, quantification from 3 mice; Video 1). Lastly, we can detect this size increase by flow cytometric analysis of AT2 cells (Fig. S2A-C; Median Forward Scatter Area quantification from 3 mice/condition), demonstrating the injury-induced phenotype is not likely an artefact of lung fixation or processing.

**Figure 1:**
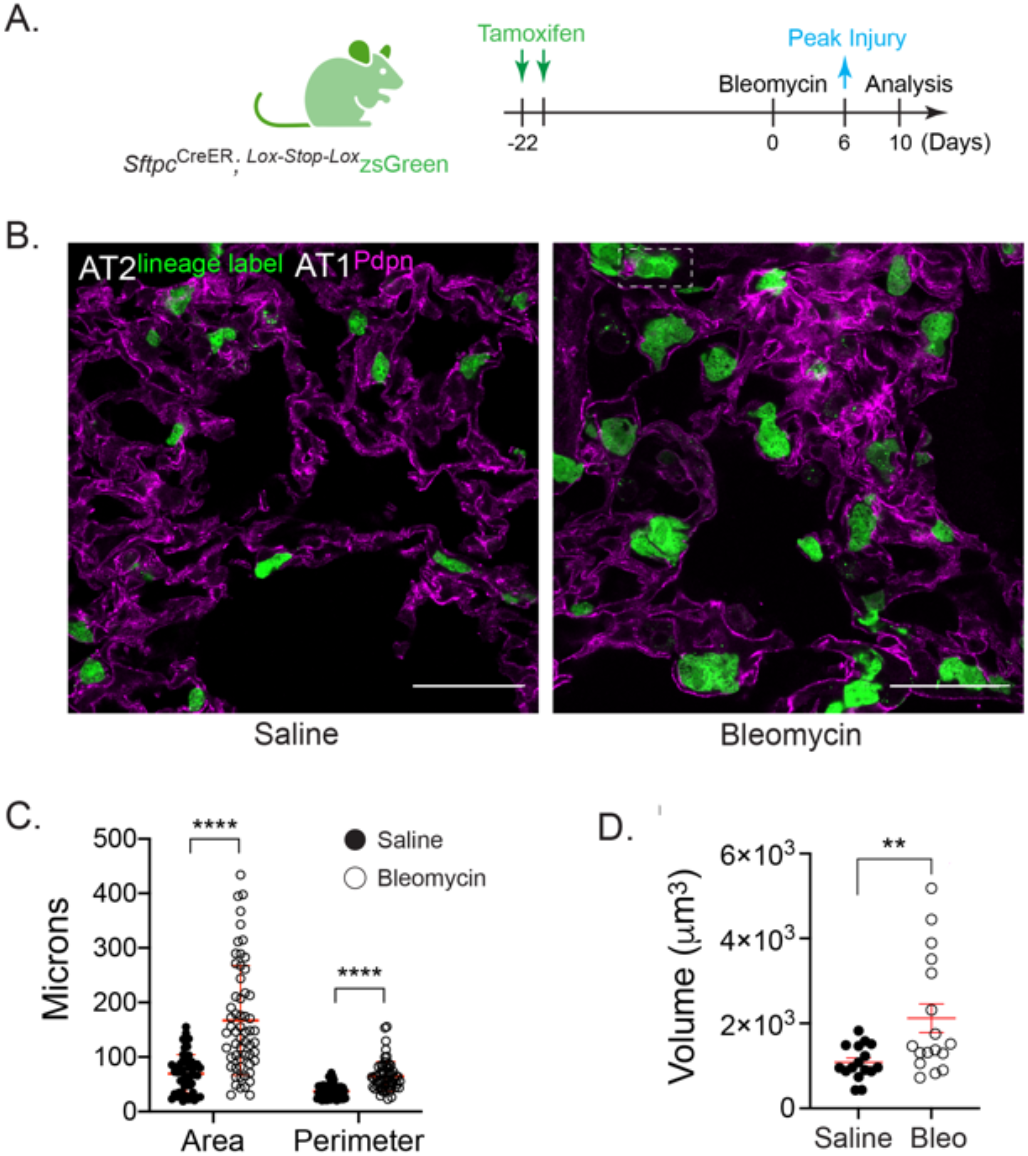
Lung injury induces AT2 cell hypertrophy. (**A**) Schematic of experimental design. Young adult *Sftpc*^CreER^; Lox-Stop-Lox^ZsGreen^ mice (3-5 months) received 10mg of tamoxifen via oral gavage 22 days prior to administration of bleomycin (0.025 units, i.t.). Lungs from saline (control) and bleomycin-treated mice were harvested on Day 10, fixed and processed for frozen sectioning and immunofluorescence analysis. (**B**) Representative lung sections showing native GFP fluorescence from AT2-lineage label and immunofluorescence staining of AT1 cells by podoplanin (Pdpn, magenta). Scale bar = 50μm. Note lineage-labeled AT2 cells appear larger after bleomycin injury. (**C**) Graph shows quantification of AT2 cell area and perimeter using image analysis in FIJI (workflow in Fig. S1A). Data points are from two mice, 5 fields of view per condition. Mean +/− SD. p < 0.0001 by t-test. (**D**) Quantification of lineage-labeled AT2 cell volumes using cleared, 200μm thick-section optical z-stack imaging (Imaris). Data points are from three saline and two bleomycin-injured mice. Mean +/− SD; p = 0.0072 by t-test. Representative fields of view in Supplemental Video 1. To ensure robustness and increase N’s, injury-induced cell size increase also validated by flow cytometry in Fig. S2.

### Lung injury promotes AT2 cell polyploidization

As cell size can scale with DNA context (26), we sought to examine nuclear features of hypertrophic AT2 cells. Interestingly, we found a number of lineage-labeled AT2 cells with two nuclei (Fig. 2A), often observing 1-2 binucleated zsGreen+ cells per field of view, estimating a rate of ~8% (Fig. 2B; see also Fig.1B inset box). To validate AT2 polyploidy via an independent method, we sought to quantify and characterize these cells by flow cytometry paired with DNA content analysis (Fig. S2A-E). Although we readily detected AT2 cells with 2N or 4N DNA content, too few 4N cells survived the cytospin procedure to reliably quantify binucleation rate differences between saline and bleomycin-treated mice (Fig. S2E). Methodological improvements will be required to reliably isolate and quantify this apparently fragile cell state, particularly given that flow cytometry is known to underestimate expansion of AT2 numbers post-injury (35).

**Figure 2:**
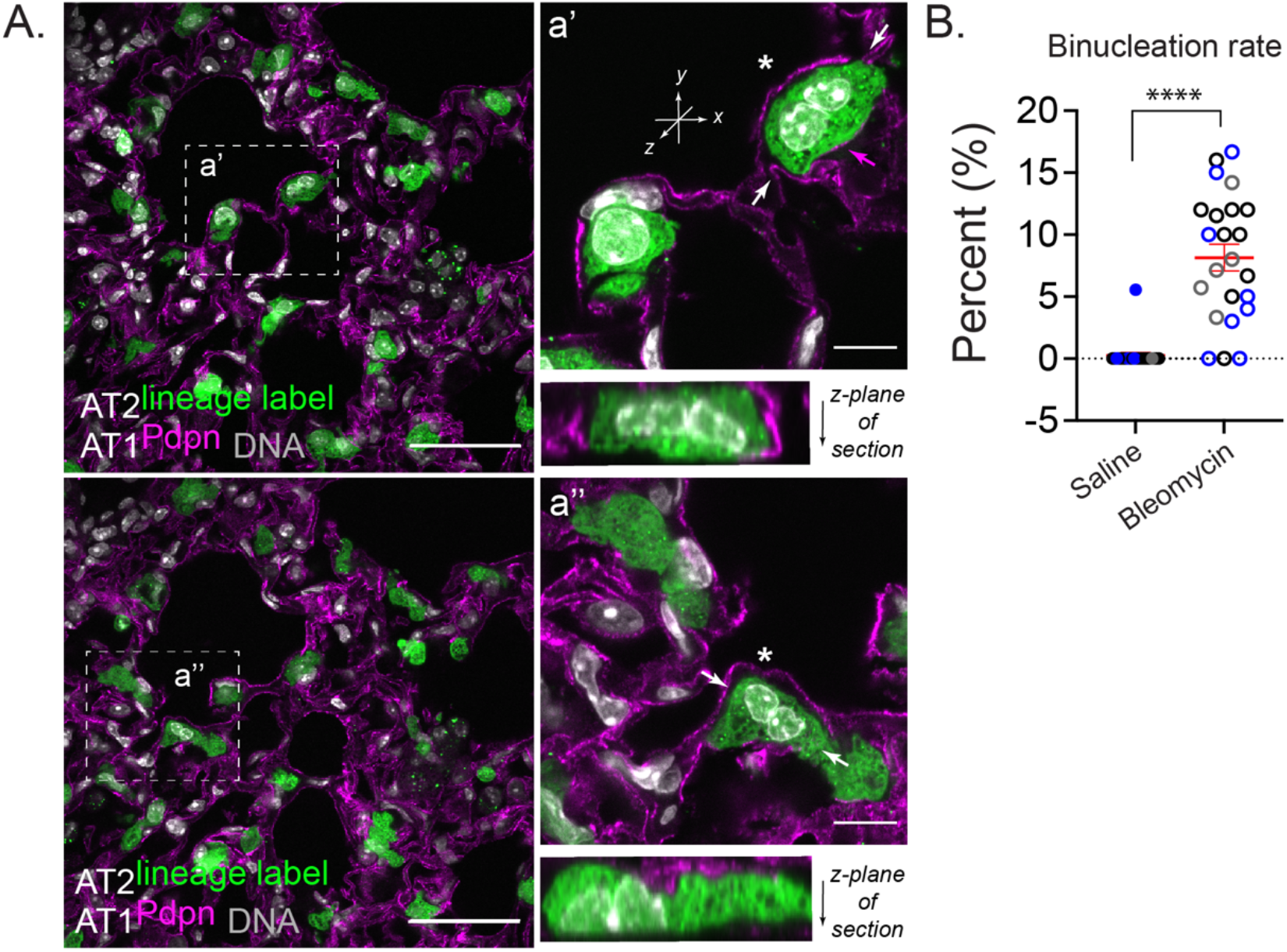
Lung injury induces polyploidization of lineage-labeled and SpC+ AT2 cells. (**A**) Schematic of experimental design as in Fig.1. Representative confocal images of lung sections from bleomycin-treated mice; two independent fields of view from low and high-magnification insets (boxed region) are shown (a’ and a”). Native GFP fluorescence from AT2-lineage label and immunofluorescence staining of AT1 cells by podoplanin (Pdpn, magenta). DNA labeling by Hoechst (gray). Scale bar = 50μm and 10 μm for inset. Asterisks show suspected binucleated AT2 cells. Arrows show plan of optical z-section, revealing that nuclei appear to touch in these cells. Magenta arrow in a’ indicates podoplanin signal. It is not possible in this image to know if podoplanin labels the binucleated AT2 or an adjacent AT1 cell (see Fig. 6D for evidence of binucleated/podoplanin positive cell). SpC+ binucleated AT2 cells are shown in Fig. S2F. (**B**) Graph shows quantification of AT2 cell binucleation. Data points are from three mice (different symbol colors/mouse), 5-8 fields of view per condition. Note the binucleation rate for one mouse (gray circles) was quantified 21 days post-bleomycin, showing persistence of binucleation through this time point. Mean +/− SD in red. p < 0.0001 by t-test. Injury-induced binucleation was also interrogated by flow cytometry in Fig. S2A-E.

Of interest, AT2 binucleation appears compatible with expression of the AT2 marker, surfactant protein C (SpC) (Fig. S2F), as well as the AT1 marker, podoplanin (Fig. 6D; *ex vivo* AT2 cell cultures). Together with recent evidence that genetically-induced AT2 binucleation is compatible with Hopx expression (37), these data suggest alveolar epithelial polyploidy may be compatible with either AT2 or AT1 cell state. To address whether AT2 polyploidy is a typical feature of the recently described AT2-to-AT1 cell transition state (21, 22), we interrogated Krt8 expression in our lung sections. While we can find large binucleated cells that highly express Krt8 protein, AT2 binucleation does not exclusively correlate with a Krt8-high state (Fig. 3). Remarkably, binucleation is a common feature of primary AT2 cultures grown on glass or filters (Fig. 4). We are confident these *ex vivo* polyploid cells derive from *bona fide* AT2 cells, as they are uniformly Krt8-positive (Fig. 4A) and can be lineage-labeled using the *Sftpc*-^CreER; Lox-stop-Lox^ ^eYFP^ system (Fig. S4A,B). Even AT2-derived organoids, which can show AT2-to-AT1 cell transition state features seen in injured lungs (23), exhibit a binucleation rate similar to *in vivo* estimates (Fig. 4B-D vs Fig. 2B). Mechanistically, live-imaging reveals binucleated cells can be generated via failed cytokinesis, further suggesting that how AT2 polyploidy is generated and maintained can be interrogated in *ex vivo* cultures (Fig. 5).

**Figure 3:**
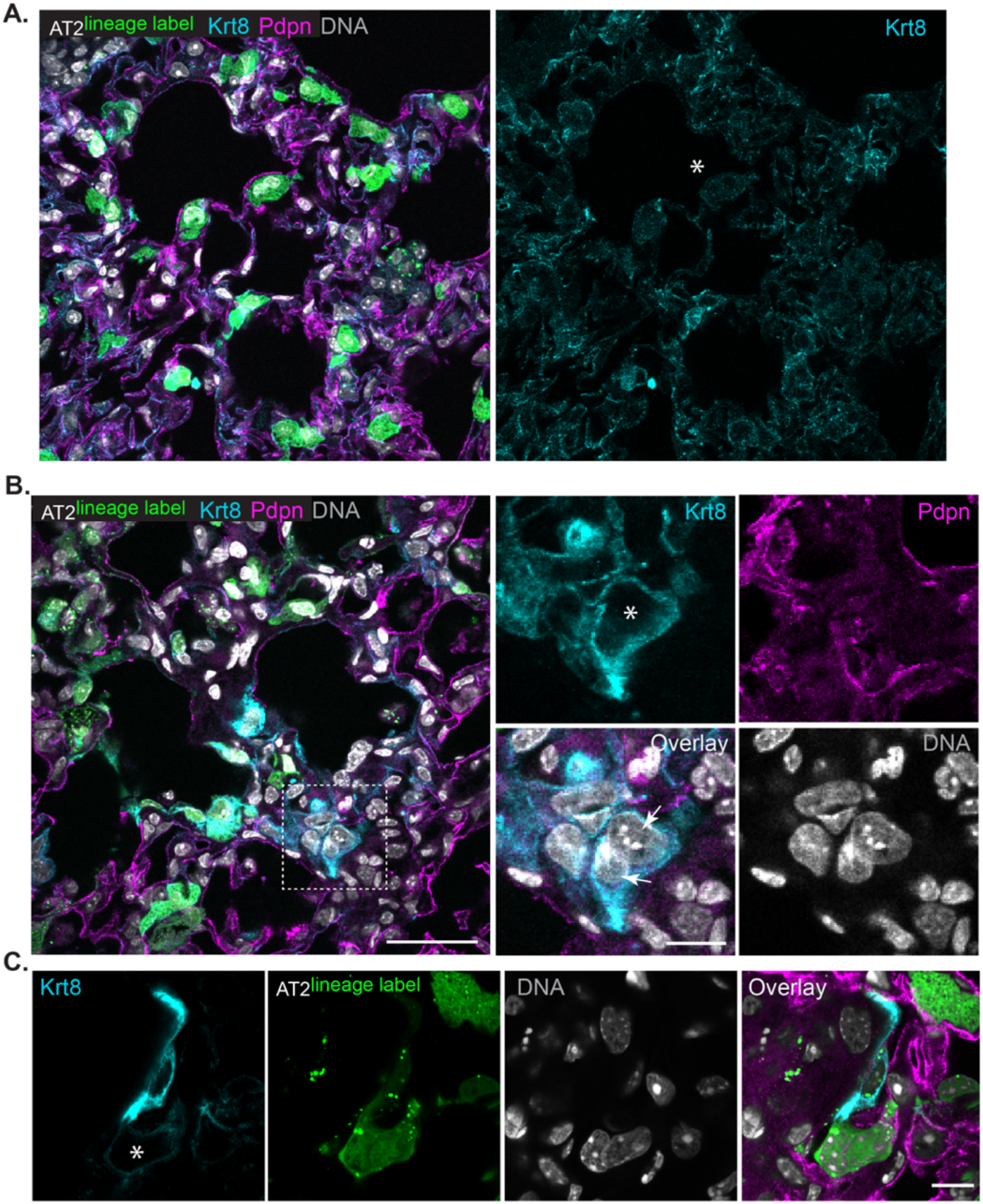
AT2 cell binucleation is not universally associated with Krt8-high expression phenotype. (**A**) Schematic of experimental design as in Fig.1. Confocal image of lung section from bleomycin-treated mice; same image as Figure 2A, now including Krt8 immunofluorescence channel (cyan). Asterisk denotes binucleated AT2 cell with normal levels of Krt8 protein. Native GFP fluorescence from AT2-lineage label and immunofluorescence staining of AT1 cells by podoplanin (Pdpn, magenta). DNA labeling by Hoechst (gray). (**B**) Confocal image of binucleated Krt8-high cell (asterisk) lacking AT2 lineage-label. Scale bar=50μm and 25 μm for inset. (**C**) Confocal image of lineage-labeled binucleated AT2 cell adjacent to lineage negative mononuclear Krt8-high cell (z-stack not shown). Scale bar=25μm.

**Figure 4:**
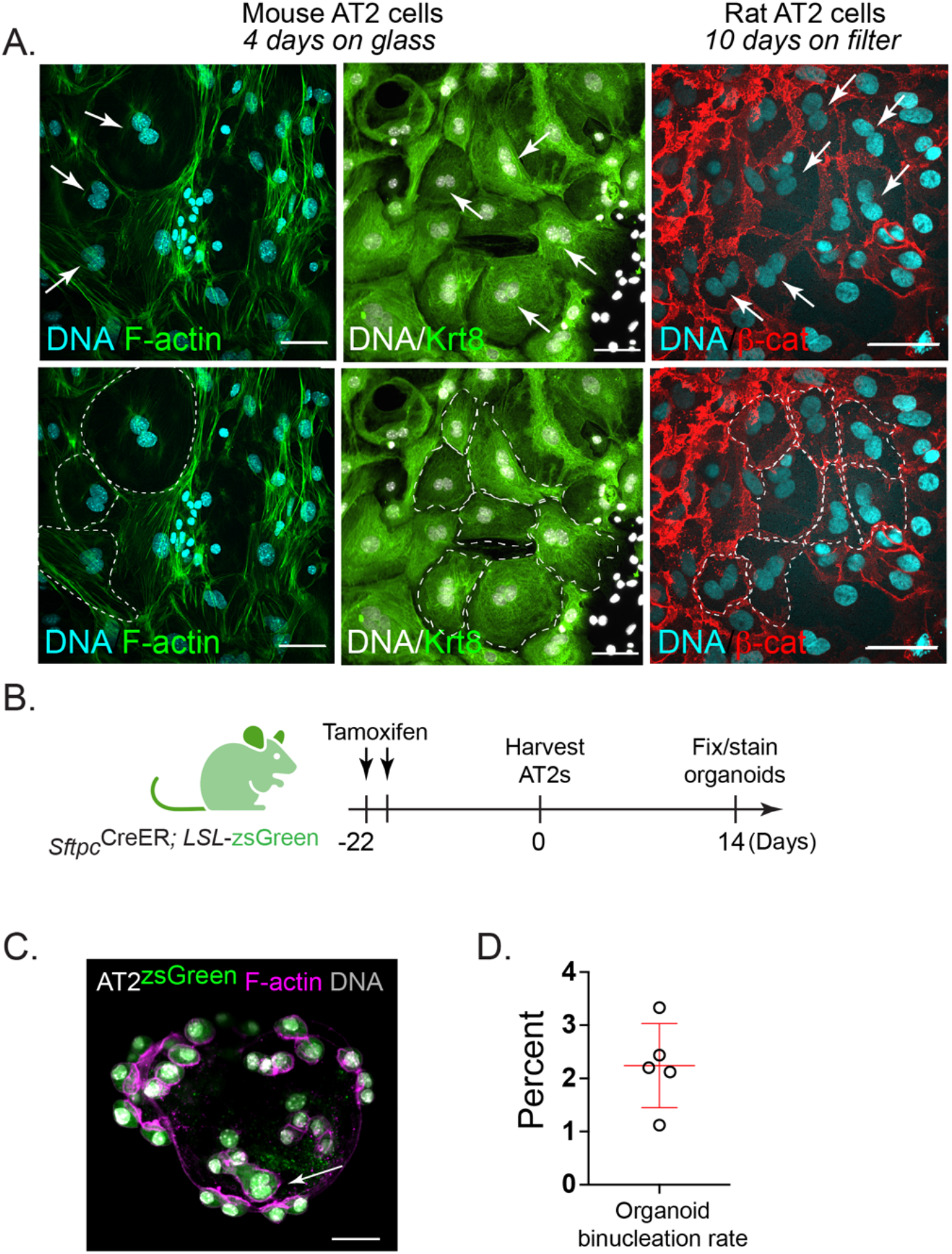
Binucleation is a feature of murine AT2 cell cultures and organoids. (**A**) Top: Confocal images of mouse AT2 cells grown 4 days on glass and rat AT2 cells grown 10 days on filters. Cells were fixed and stained with Hoechst (DNA in cyan or gray), phalloidin (F-actin, green), Krt8 (green) or β-catenin (red) as indicated. Top: Arrows (white) indicate two nuclei “kissing” without an intervening junction (by F-actin or β-catenin staining). Bottom: Dotted lines (white) outline binucleated cells. (**B**) Schematic for activation of AT2-lineage label before AT2 isolation for organoid growth. (**C**) AT2 cell (green), F-actin (magenta) and DNA (gray) with lineage-labeled binucleated cell (arrow). (**D**) Rate of AT2 binucleation in organoids. Scale bars=50μm.

**Figure 5:**
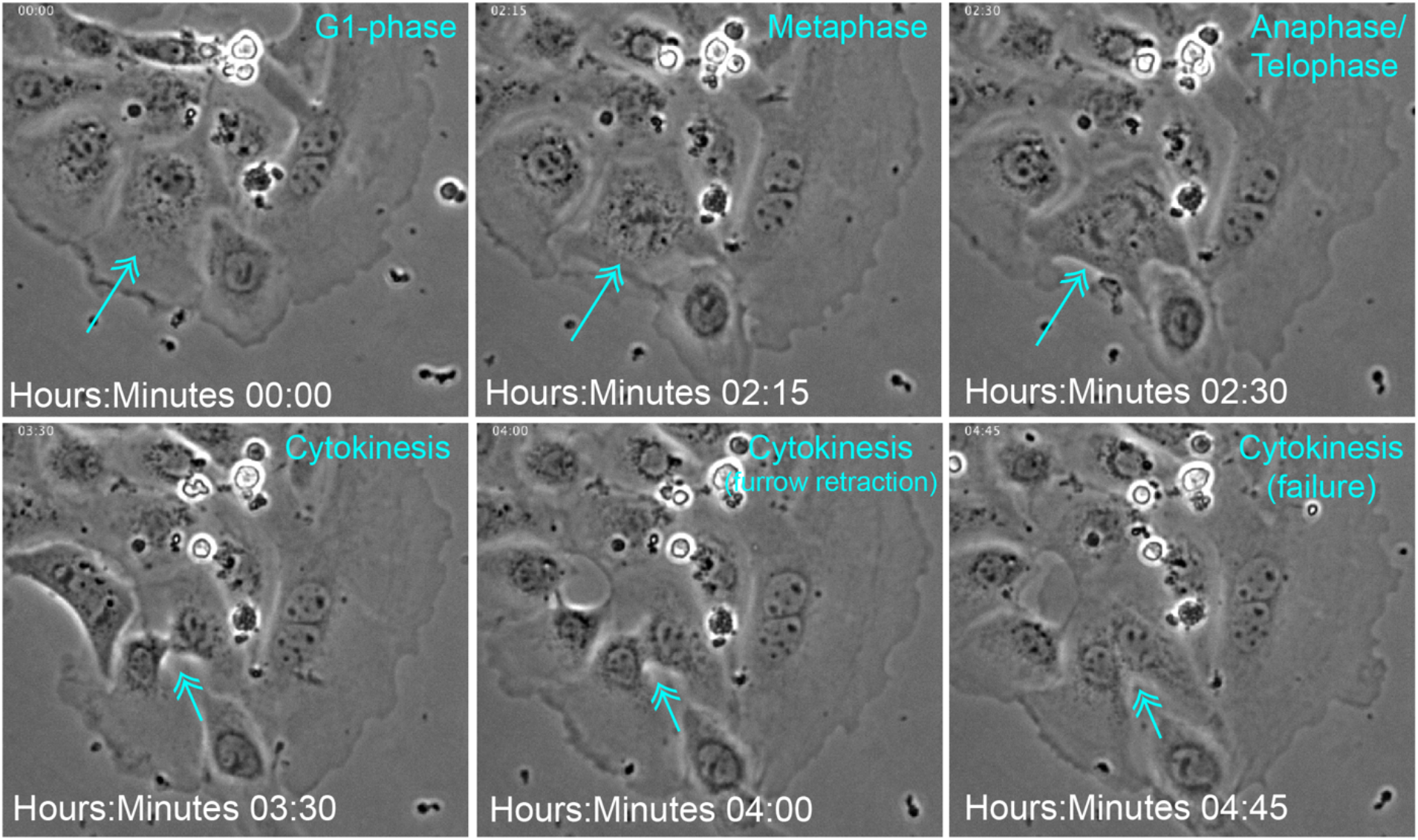
AT2 cell binucleate via failed cytokinesis. Mouse AT2 cells were isolated and phase-contrast live-imaged every 15-minutes (Methods). Time stamp and evident cell-cycle phases are shown in cyan. Arrow indicated dividing cell.

**Figure 6:**
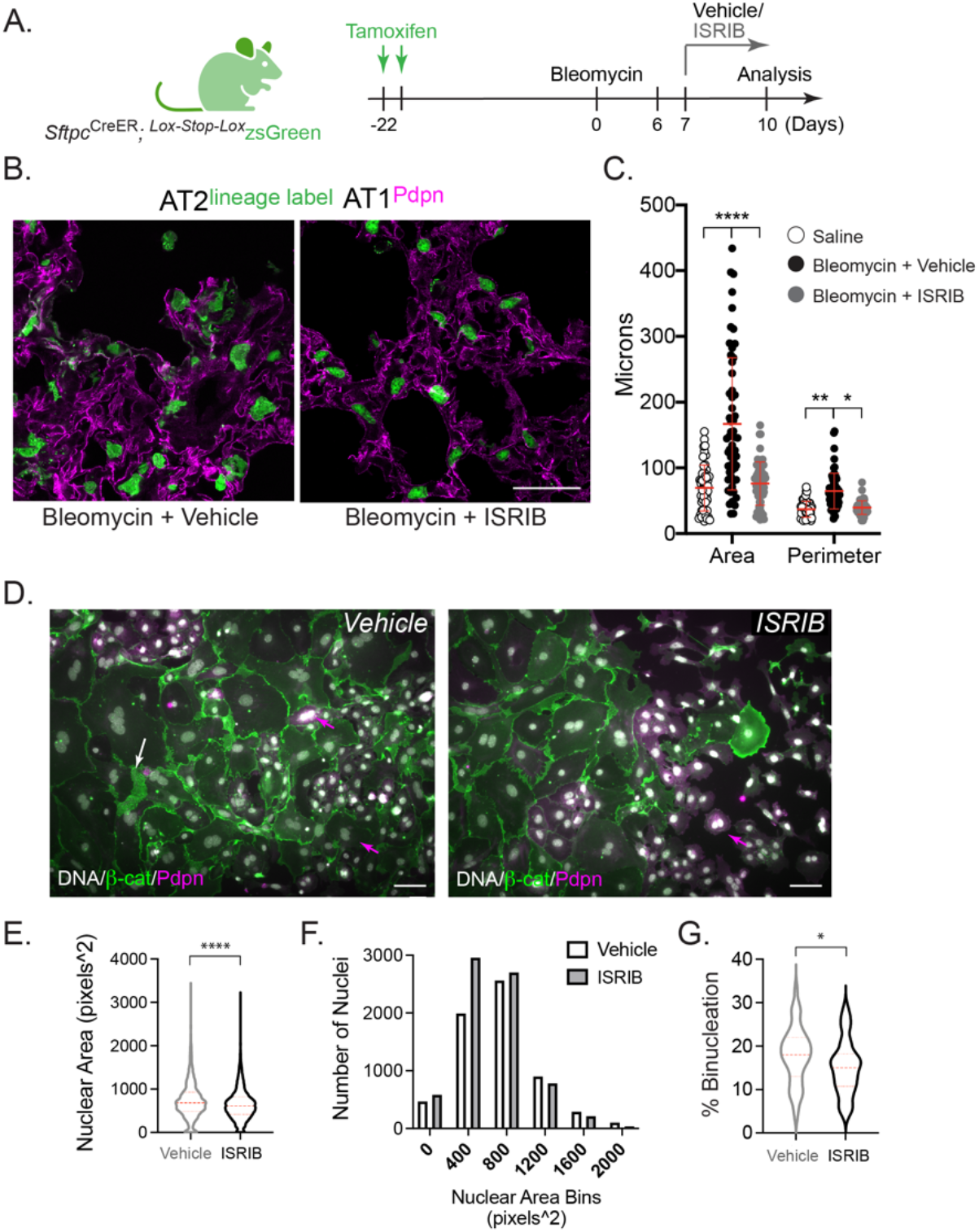
ISRIB limits AT2 hypertrophy and polyploidy cell-autonomously. (**A**) Schematic for AT2 lineage-trace before bleomycin-injury with and without ISRIB therapy. (**B**) Confocal images of fixed frozen lung sections with native zsGreen and immunofluorescence of AT1 marker, podoplanin. Scale 50μm. (**C**) Scatter plot of zsGreen object sizes by FIJI as described in Figure S1. Graph identical to Fig.1C with ISRIB data added. Significance by ANOVA. (**D**) Epifluorescence overlay images of mouse AT2 cell Day2-cultures incubated with Vehicle/ISRIB (0.025μm) through Days 4-5. Arrows (magenta) show podoplanin+ binucleated cells that are peripheral to large flat bi-/tri-nucleated cells with diminishing podoplanin and high levels of β-catenin (white arrow). Scale 50μm. (**E**) Nuclear area (DNA) measurements by ImageJ as proxy for cell size allowed for unbiased analysis. N=6829 (Vehicle) and 7387 (ISRIB) nuclei measured. Data averaged from 3 independent AT2 isolations for cells quantified at 48 and 72-hours +/-ISRIB. p < 0.0001**** by Mann-Whitney test. (**F**) Histogram of data in **E**. (**G**) Binucleation events quantified as from 44 (Vehicle) and 43 (ISRIB) fields of view. p=0.035* by Mann-Whitney test. Dotted red lines reflect median, first and third quartiles.

Given limitations of assessing polyploidy from frozen or thick sectioning (Fig. S1B), together with evidence that mononuclear polyploidy also exists (38), we sought an independent method to assess the polyploidy rate in injured AT2 cells. A recent study showed that the stochastic, multi-color Confetti reporter system can be used to assess mononuclear polyploidy in mouse tissues, notably liver (39). While the Confetti reporter mouse is better known for tracing clonal cell relationships in development and injury (14, 40), this tool can also validate polyploid cells in tissues with single cells expressing two or more fluorescent proteins (Fig. S3A). In contrast to cell clone tracing, the lineage label must be activated by tamoxifen *after genome duplication* to detect single cell polyploidy. By subjecting *Sftpc*^Cre-ER^ x *Rosa*^Confetti^ mice to the bleomycin model of injury, we could identify large, partially flattened cells that express both green fluorescent protein (GFP) and red fluorescent protein (RFP) lineage labels surrounding what appears as a single nucleus of lineage-label exclusion (Fig. S3B,C). We also identified small round, GFP/RFP-dual-positive cells that likely reflect lineage-labeled AT2 cells which have doubled their DNA content and arrested at the G2-phase of the cell cycle (Fig. S3D). As these dual-positive cells appear more rare than the AT2 binucleation rate estimated in Figure 1 (e.g., < 2% lineage-labelled AT2 cells), we may be missing polyploidization events during the AT2-to-AT1 transition, when *Sftpc*^Cre^-expression is likely waning. Crossing the *Rosa*^Confetti^-reporter strain to a Cre-driver whose expression increases during the AT2-to-AT1 transition may be required to assess the full extent of alveolar epithelial polyploidization during the lung injury-repair sequence. Nonetheless, these data show that lung injury can generate binucleated polyploid AT2 cells (Fig. 1), and possibly polyploid mononuclear AT2 cells (Fig. S3C), where polyploidization likely occurs during the post-injury, reparative phase of the bleomycin model.

### Inhibiting the Integrated Stress Response attenuates AT2 hypertrophy

To determine whether injury-induced AT2 hypertrophy and polyploidy can be inhibited or reversed, we considered small molecule approaches known to target stress-activated pathways, recently suggested to guide injured AT2 cells through their vulnerable transition towards an AT1 fate (23, 24). The integrated stress response (ISR) is a highly conserved signaling network that helps cells adapt to a range of environmental stresses, through activating kinases that inhibit bulk protein translation in favor of the translation of select mRNAs encoding transcription factors such as ATF4 that promote the transcription of molecular chaperones and other stress adaptive genes (41). A central integrator of these diverse stress-responsive kinases is the eIF2 complex, where phosphorylation of eIF2α at a single serine residue limits translation (42). In this way, eIF2α phosphorylation and its diverse downstream consequences comprise the ISR (43, 44). A small molecule integrated stress response inhibitor, ISRIB, was found to render cells insensitive to eIF2α phosphorylation, thereby restoring translation capacity to normalize cell function (45–47) and limit a range of tissue pathologies in murine disease models (48–51). Our team recently discovered that ISRIB reduces fibrosis in mouse models (24). Since ISRIB therapy limited abundance of lineage-labelled AT2 cells co-expressing KRT8, yet increased abundance of lineage-labeled cells expressing podoplanin, we reason that ISRIB may attenuate fibrosis by limiting abundance of AT2 cells stalled at the AT2-to-AT1 transition state, favoring their full transition to an AT1 fate (24). Indeed, evidence that bleomycin-injured AT2 cells show activation of ISR-associated genes during this transdifferentiation step (24), further suggests that persistent activation of the ISR in AT2 cells can thwart alveolar epithelial repair. However, the particular morphogenetic features of the AT2-to-AT1 transition that drive persistence of this stress state remain unclear. We find that bleomycin-injured *Sftpc*^Cre-ER; Lox-Stop-Lox^zsGreen mice treated with ISRIB show complete inhibition of AT2 hypertrophy and no evidence of binucleation (Fig. 6A,B). Although ISRIB did not prevent AT2 cell binucleation in *ex vivo* cell cultures, ISRIB treatment did limit the rate of binucleation, as well as formation of the largest cells, inferred from nuclear size within these cells (Fig. 6C-G). These data suggest that injury-induced AT2 hypertrophy and polyploidy are sensitive to inhibition of the ISR. Evidence for modest cell size attenuation in *ex vivo* AT2 cultures suggests that ISRIB may limit hypertrophic growth through an AT2 cell-autonomous mechanism.

## Discussion

For decades, clinical pathologists have described the development of “hypertrophic” AT2 cells after injury, but mechanisms that drive this state and degree to which it benefits and/or constrains the lung injury-repair cycle is not known. We show that some of this hypertrophic growth is associated with AT2 polyploidy, most notably binucleated AT2 cells. *Ex vivo* analysis suggests the route to polyploidy is via failed cytokinesis during the AT2-to-AT1 flattening process. Remarkably, attenuating the ISR with ISRIB limits the abundance of hypertrophic, polyploid AT2 cells after bleomycin injury and *ex vivo* culture, suggesting the effects of ISRIB may be AT2 cell-autonomous. Given recent high resolution time-series sampling of the injured lung and analysis by single cell RNA-sequencing revealing that the AT2-to-AT1 transition state expresses genes associated with the ISR (22), we suggest that lung injury persistently activates the ISR in AT2 cells as they divide, enlarge and flatten to repair the alveolar epithelium, increasing their susceptibility to cytokinesis failure into polyploid AT2 cells. A key consequence of AT2 polyploidization will be loss of the AT2 stem cell daughter, which may compromise future regenerative potential of alveoli served by that AT2 progenitor. We propose that attenuating the ISR helps guide AT2 cells through this vulnerable morphogenetic sequence of cell-growth, division and trans-differentiation without polyploidy (Fig. 7, Model).

**Figure 7:**
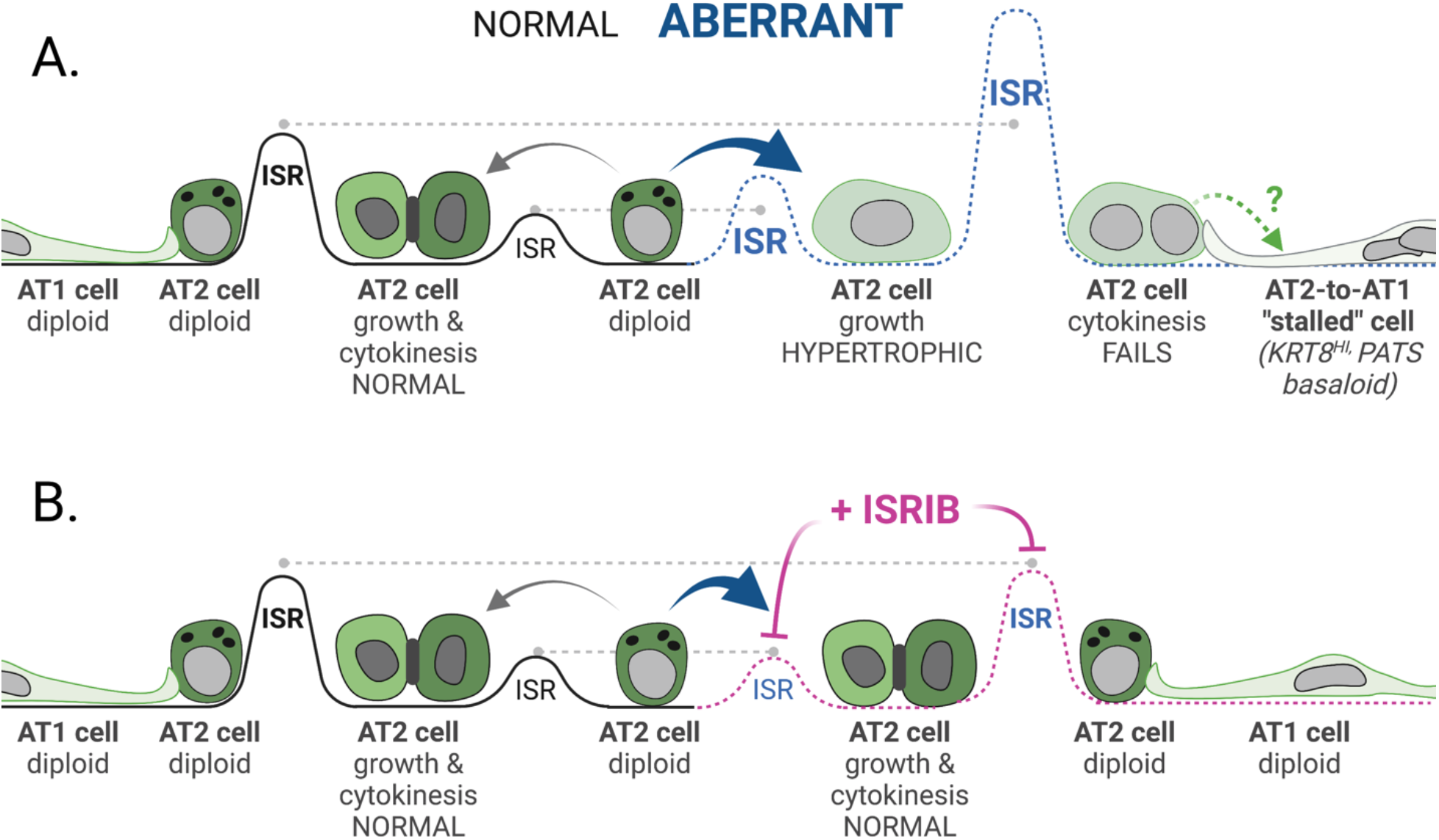
Model for alveolar epithelial stress-induced hypertrophy and polyploidization in lung injury and repair. (**A**) AT2 “aberrant” repair (dotted blue line) due to persistent ISR, AT2 hypertrophic growth, cytokinesis failure and stalled AT2-to-AT1 cells (right). (**B**) ISRIB therapy (dotted pink line) restores ISR to levels for normal AT2-to-AT1 growth, division and differentiation (dotted gray line). Schematic with BioRender software.

Evidence that AT2 binucleation does not exclusively correlate with cells expressing the highest levels of KRT8 protein is noteworthy. While elevated *Krt8* expression is associated with an aberrant AT2-to-AT1 transition state (21, 22), perhaps this transition is only one of a number of transcriptionally distinct routes cells take as they repair the injured alveolar epithelium. Alternatively, the Krt8-high/*Sftpc*^*CreER; zsGreen*^-negative binucleated cells detected in Figure 3B suggests alveolar epithelial polyploidization may also occur via *Sftpc*-lineage negative progenitors (52). Given the increased size and presumed fragility of polyploid alveolar cells, we suspect they are inadequately represented in current lung single cell RNA-sequencing pipelines. To understand how these cells specifically contribute to alveolar repair, it will be necessary to revise existing cell sorting pipelines to quantify the abundance and degree of alveolar epithelial polyploidy, as well as isolate these cells for downstream transcriptomic analyses to establish the distinct features of the polyploid cell state (33).

Recent injury-repair studies using live-imaging of zebrafish epicardium as a simple epithelial model system reveal that cells along a closing wound-front manifest higher tension and rates of binucleation due to failed cytokinesis (28). While one study suggests this may be due to excessive integrin-based extracellular matrix adhesive tension that ultimately interferes with contractility of the actin-myosin cytokinetic ring (53), there are many routes to cytokinesis failure caused by reduced or elevated Rho-signaling (54). Endoplasmic reticulum (ER)-stress itself can lead to cytokinesis failure (55, 56). Moreover, PKR, a dsRNA-dependent Protein Kinase and upstream activator of the ISR, can be activated by mitosis onset, where nuclear membrane breakdown releases endogenous dsRNAs that bind and activate PKR (57). This activation is required for cytokinetic fidelity, underscoring the relationship between ISR signaling and cell division. Future work will be required to determine whether cytokinesis is more vulnerable to ATF4-dependent versus independent arms of the ISR.

Although bleomycin-injury leads to binucleation in only ~5% of lineage-labeled AT2 cells, total alveolar epithelial cell polyploidy may be much greater, particularly given evidence for mononuclear polyploidy across tissues (38). Newer imaging methods that allow simultaneous assessment of DNA content and cell cycle phase will be required to distinguish true mononuclear polyploidy (i.e., ≥4N DNA content in a G_1_-phase cell) from AT2 cells arrested post-genome doubling at the G_2_-phase (i.e., also 4N DNA content) (58). Another limitation of this study is our reliance on the DNA-damaging agent bleomycin as an injury model, which could bias the alveolar repair response towards AT2 polyploidy. Future work will be required to determine the generality of this paradigm, and whether AT2 polyploidy can be induced by other forms of injury (e.g., viral or pneumonectomy model of alveologenesis).

Conceptually, it will be important address whether AT2 polyploidization is restricted to the transdifferentiating daughter, sparing normal ploidy of the renewed AT2 or if lung injury promotes AT2 endocycling/endomitosis, resulting in a polyploid facultative progenitor. The former would generate polyploid AT1 cells, which might allow AT1 cells to become larger with excess DNA copies for environmental adaptation (25). Indeed, a recent proof-of-concept study, which genetically induced cytokinesis failure in AT2 cells, found that binucleation does not obviously prevent trans-differentiation towards at AT1 fate (37). These data are consistent with our own evidence that binucleation is compatible with expression of the AT1-marker, podoplanin (Fig. 6D). Polyploidization of AT2 cells could provide extra DNA copies for surfactant biosynthesis, similar to how mammary epithelial cells become polyploid during lactation to maximize milk production (30). However, the induction of AT2 polyploidy would likely compromise its progenitor function, which requires faithful segregation of additional chromosomes during mitosis for stem cell renewal (38). Even heart tissue, which comprises large binucleated cardiomyocytes suited to contractility, relies on a low percentage of mononuclear diploid progenitor cells (2-14%) to drive regeneration post-injury. Mouse strains with a greater percentage of these mononuclear diploid cells (14%) recover from injury faster than strains with a lower percentage (2%) (59). Given this precedence, we reason that even a small percentage of AT2 cells becoming polyploid during an injury event (~5% per injury) could ultimately contribute to substantial loss of AT2 progenitor renewal over a lifetime, absent alternate mechanisms for AT2 progenitor replacement, such as via *Sftpc*-negative progenitors (60).

It will be important to understand how alveolar epithelial cells sustain the polyploid state. Forced-induction of polyploidy in retinal pigmented epithelial cells requires activation of p53 signaling (61). Given recent evidence that forced-activation of p53 may facilitate the AT2-to-AT1 transition during lung injury (23), it is possible that induced polyploidy plays a beneficial role in the earliest stages of alveolar epithelial repair. However, evidence that the integrated stress response inhibitor, ISRIB, attenuates formation or duration of hypertrophic alveolar epithelial cells (this study) and promotes lung repair in the bleomycin model (24), raises the possibility that maintaining polyploid cells for too long may be detrimental.

AT2 hypertrophic growth not associated with polyploidy may also be problematic for the injury-repair sequence. Young cells manipulated to grow to a large size, even without a corresponding increase in DNA content (i.e., polyploidy), can manifest phenotypes observed in senescent cells (62, 63). Conversely, older cells are associated with volume increase, raising the possibility that cytoplasmic dilution of key factors may contribute to senescence-associated phenotypes (62). Indeed, this same research team found that cell size is a determinant of stem cell (hematopoietic) potential during aging, where cell enlargement contributes to functional decline (64). Thus, hypertrophic growth independent of polyploidy can be maladaptive. Since most of our lineage labeled AT2 cells show size increase post-injury, whereas the binucleation rate is only ~5%, absent substantially higher levels of mononuclear polyploidy, it is likely that AT2 hypertrophy and ISR activation are upstream of and causal to the polyploidization step (Fig. 7). We speculate, therefore, that an inappropriately timed hypertrophic growth spurt may interfere with successful completion of mitosis. We hypothesize that ISRIB limits hypertrophic growth (and polyploidy) through an ATF4-dependent or independent (i.e., bulk-translation) mechanism, which can be addressed using existing ATF4 gain- and loss-of-function mice. Ultimately, adaptive or maladaptive functions of hypertrophic normal (2N) versus polyploid (≥4N) AT2 cells will be informed by transcriptomic studies that consider DNA content as a variable. The lung needs to repair in a manner that does not massively increase cell numbers (65). Epithelial polyploidization post-injury may be a way to repair such tissues where excessive cellularity may be problematic. Epithelial polyploidization consequent to injury is a process with short-term benefits (e.g., larger cells with enhanced barrier function/reduced leak, increased DNA copies for adaptation to stressful environments), but long-term consequences (e.g., reduced stem cell maintenance due to loss of self-renewed mononuclear AT2). Our data raise the possibility that modulators of the ISR may be leveraged to maintain AT2 progenitor function during injury. Future work will be required to understand the means by which ISRIB limits formation of or guides the transition of hypertrophic, polyploid AT2 cells towards an AT1 phenotype, and degree to which induced polyploidy contributes to alveolar epithelial cell heterogeneity or adaptation in lung disease.

## Materials and Methods

Key resources Table

Key resources Table
| REAGENT or RESOURCE | SOURCE | IDENTIFIER |
| --- | --- | --- |
| **Antibodies** |  |  |
| Rat anti-mouse I-A/I-E (MHC II), BUV395 | BD Biosciences | Cat#743876, RRID:AB_2741827 |
| Rat anti-mouse CD45, APC | BioLegend | Cat#102410 |
| Rat anti-mouse CD31, APC | BioLegend | Cat#103112 |
| Rat anti-mouse CD326 (Ep-CAM), PE/Cy7 | BioLegend | Cat#118216 |
| Rat anti-mouse CD11c, PE/Cy7 | BioLegend | Cat#117318 |
| Propidium Iodide | ThermoFisher Scientific | Cat#P1304MP |
| Biotin Rat Anti-mouse CD16/32 | BD Biosciences | Cat#553143 |
| Biotin Rat Anti-mouse TER119/Erythroid Cells | BD Biosciences | Cat#553672 |
| Biotin Rat Anti-mouse CD45 | BD Biosciences | Cat#553078 |
| Biotin Rat Anti-mouse CD90 | BD Biosciences | Cat#554896 |
| Anti-Prosurfactant Protein C (proSP-C) | Millipore Sigma | Cat#AB3786 |
| Anti-β-catenin | Millipore Sigma | Cat#06-734 |
| Anti-β-catenin | BD Biosciences | Cat#610153 |
| Podoplanin | DSHB (University of Iowa) | Cat#8.1.1, RRID:AB_531893 |
| Krt8 | DSHB (University of Iowa) | Cat#TROMA-I, RRID:AB_531826 |
| Alexa Fluro 680 Phalloidin | Invitrogen | Cat#A22286 |
| Goat anti-rabbit Alexa Fluor 488 | Invitrogen | Cat#A11008 |
| Goat anti-mouse Alexa Fluor 568 | Invitrogen | Cat#A11004 |
| Goat anti-hamster Alexa Fluor 594 | Invitrogen | Cat#A21113 |
| Goat anti-rat Alexa Fluor 546 | Invitrogen | Cat#A11081 |
| **Chemicals, Peptides, and Recombinant Proteins** |  |  |
| Bleomycin | Fresenius Kabi | Cat#0703-3154-01 |
| 0.9% saline chloride | BBraun | Cat#R5201-01 |
| ISRIB | AdooQ BioScience | Cat#A14302 |
| Tamoxifen | Sigma | Cat#T5648 |
| Corn oil | Sigma | Cat#C8267 |
| Vybrant DyeCycle Violet Stain | Invitrogen | Cat#V35003 |
| Hoechst 33342 | ThermoFisher Scientific | Cat#62249 |
| Draq5 | ThermoFisher Scientific | Cat#62251 |
| ROCK1 inhibitor Y-27632 | Selleckhem | S1049 |
| ProLong Gold Antifade Mountant | ThermoFisher Scientific | Cat#P36934 |
| Eriochrome black | Sigma | Cat#E2377 |
| CD326 (EpCAM) Microbeads, mouse | Miltenyi Biotec | Cat#130-105-958 |
| Matrigel | Corning | Cat#356231 |
| Collagenase | Millipore Sigma | Cat#C0130 |
| Elastase | Worthington | Cat#LS002292 |
| Deoxyribonuclease (DNase) I | Sigma | Cat#D4513 |
| Dispase | Corning | Cat#47743-724 |
| PharmLyse (Red Blood Cell Lysis) | BD Biosciences | Cat#555899 |
| Purified Rat Anti-mouse CD16/CD32 (Mouse FcBlock) | BD Biosciences | Cat#553142 |
| Streptavidin MagneSphere Paramagnetic Particles | Promega | Cat#Z548C |
| Mouse CD45 MicroBeads | Miltenyi Biotec | Cat#130-052-301 |
| autoMACS Running Buffer – MACS Separation | Miltenyi Biotec | Cat#130-091-221 |
| Critical Commercial Assays |  |  |
| SHIELD | LifeCanvas Technologies | Cat#EI-Z1001,#SH-BS,#SH-ES,#SH-ON |
| MultiMACS Cell24 Separator | Miltenyi Biotec |  |
| Experimental Models: Organisms/Strains |  |  |
| Mouse: C57BL/6J (8-12 weeks old) | The Jackson Laboratory | JAX: 000664 |
| Mouse: *Sftpc*<sup>CreER</sup> | The Jackson Laboratory | JAX: 020940 |
| Mouse: R26R<sup>ZsGreen</sup> | The Jackson Laboratory | JAX: 007906 |
| Mouse: *Sftpc*<sup>CreER</sup> | International Mouse Strain Resource | MGI: 5444645 |
| Mouse: R26R<sup>eYFP</sup> | The Jackson Laboratory | JAX: 006148 |
| Mouse: R26R<sup>Confetti</sup> | The Jackson Laboratory | JAX: 013731 |
| Software and Algorithms |  |  |
| FIJI/ImageJ (version: 2.1.0/1.53c) | Schneider, CA et al., 2012 | https://imagej.nih.gov/ij/ |
| Stardist plugin | Schmit, U et al., 2018 | Cell/nuclear size quantification<br>https://arxiv.org/abs/1806.03535 |
| Imaris |  | https://imaris.oxinst.com/ |
| Other |  |  |
| Vibratome | Leica Microsystems | VT1200s |
| Cryostat | Reichert-Jung | Cryocut 1800 |
| W1 Spinning Disk Confocal | Nikon Instruments |  |
| BioStation CT or other | Nikon Instruments |  |
| Axioplan2 epifluorescence microscope | Zeiss; 20x objective (Air) | AxioCAM HR Camera with AxioVision 4.8 software |
| A1R Confocal | Nikon Instruments |  |

### Methods Details

#### Mice

C57BL/6 mice were obtained from Jackson laboratories (Jax stock 000664). *Sftpc*^CreER^ (52), Ai6-R26^zsGreen^, *Sftpc*^CreERT2;R26ReYFP^ (kindly provided by Ed Morrisey, University of Pennsylvania)(66, 67) and *Sftpc*^CreER;R26R-Confetti^ (14) (Jax stocks 028054, 007906, 006148, 013731). Only male mice were used in bleomycin-injury experiments based on established injury severity in males relative to females, which improves dynamic range of assays for comparisons (68). Eight to twelve week old mice were used as young adult mice. All experimental protocols were approved by the Institutional Animal Care and Use Committee at Northwestern University (Chicago, IL, USA). All strains including wildtype mice are bred and housed at a barrier- and pathogen-free facility at the Center for Comparative Medicine at Northwestern University.

### Cre recombination, bleomycin administration, ISRIB administration

For nuclear activation of Cre-ER, tamoxifen was dissolved in sterile corn oil at 10 mg/100μL and administered via oral gavage once a day for two days to induce Cre-recombination of floxed alleles for lineage-tracing. For bleomycin injury, mice were anesthetized and intubated prior to the intratracheal administration of bleomycin (0.025 units in 50μL 0.9% saline). Lungs were harvested at indicated time points for downstream analyses (e.g., fixation and frozen sectioning for immunofluorescence analysis, AT2 cell isolation by negative selection (69) or EpCam+ selection (70) and flow cytometry methods (24, 71) with modifications (below). ISRIB (AdooQ BioScience, A14302) was first reconstituted in DMSO to the concentration 4mg/mL and then further diluted using PBS to a final concentration of 0.3125 mg/mL (72). ISRIB was prepared at 2.5 mg/kg and administered daily through intraperitoneal injection in a volume of 200μL. Control animals were treated with vehicle (DMSO/PBS) alone.

#### Tissue preparation, AT2 isolation, flow cytometry and sorting

Tissue preparation for AT2 isolation and analyses was performed as described (24, 71), with modifications. Briefly, mice were euthanized and lungs were perfused through the right ventricle with 10ml of HBSS. Lungs were removed and infiltrated with dispase and 0.2 mg/ml DNase I dissolved in HBSS with Ca^2+^ and Mg^2+^, using syringe with 30G needle. Lungs were chopped with scissors and incubated in DMEM containing 5% FBS for 45 minutes with agitation. The tissue suspension was filtered through a 70 μM strainer and washed in 10 mL media (DMEM; 5% FBS). The pellet was then suspended in red blood cell (RBC) lysis buffer for 1 minute, washed with media, and filtered through 40 μM strainer. The cells were incubated with EpCAM magnetic beads and AT2 were collected using the MultiMACS Cell24 Separator (Miltenyi Biotec). Automated cell counting was performed using Nexcelom K2 Cellometer with AO/PI reagent. Cells were incubated with FcBlock (BD Biosciences, San Jose, California), and stained with a mixture of fluorochrome-conjugated antibodies, as listed above. DNA content was quantified using Vybrant DyeCycle Violet. Single color controls were prepared using BD CompBeads (BD Biosciences, San Jose, California) and Arc beads (Invitrogen, Waltham, Massachusetts). Flow cytometry and cell sorting were performed at the Northwestern University Robert H. Lurie Comprehensive Cancer Center Flow Cytometry Core facility (Chicago, Illinois). Data were acquired on a custom BD FACSymphony instrument using BD FACSDiva software (BD Biosciences, San Jose, California). Compensation and analysis were performed using FlowJo software (TreeStar). Each cell population was identified using sequential gating strategy. The percentage of cells in the live/singlets gate was multiplied by the number of live cells using Cellometer K2 Image cytometer to obtain cell counts. Cell sorting was performed using BD FACSAria III instrument using 100μm nozzle and 40 psi pressure. *Cytospin*: Following AT2 sorting, collected cells were spun at 300g for 10 minutes. Following resuspension in PBS, cells were plated onto slides. Slides were spun in cytospin filters at 100g for 5 minutes followed by staining with phalloidin (1:50) and Hoescht (1:10,000).

#### AT2 isolation for ex vivo two-dimensional culture

Lungs were inflated intratracheally with Dispase and incubated for 45 minutes with agitation. Lungs were chopped and incubated in 7mL of media containing 10% FBS and DNase I (1 mg/mL) for 10 minutes. Tissue suspensions were filtered through a 70μM strainer. The cell pellets were resuspended in RBC lysis buffer for 2 minutes. Alveolar epithelial cells were enriched through negative selection against biotinylated CD16/32, CD45, CD90, and TER119 antibodies. The cell suspensions were then incubated at 37°C for 2 hours followed by gentle panning. Cells were counted and cultured in complete media (10% FBS, 2% Penicillin and Streptomycin) onto 12mm glass coverslips. Starting 48 hours after isolation, cells received a daily media change either containing 0.025μM ISRIB or DMSO. Cell samples were pooled from 3 mice. Rat AT2 cells were collected as previously described (73). AT2 cell purity was validated by cytospin analysis (as in Fig. S2), with staining for surfactant protein C, keratin 8 and vimentin; Vimentin+ cells are less than 3% and surfactant protein C is expressed in ~95% of the keratin 8 population.

#### AT2 organoid culture

For organoid culture, cells were prepared as previously described (14) with modifications. Mice received tamoxifen 14 days prior to isolation. AT2 isolation was conducted as described above. For fibroblast isolation, lungs were inflated with 1 mL of collagenase (200 U/mL), elastase (4 U/mL), and DNase solution (0.25 mg/mL). Lungs were cut into pieces and incubated for 30 minutes at 37°C in digestion buffer. Lungs were processed in C-tubes (Miltenyi Biotec, 130-093-237) using GentleMACS/OctoMACS dissociator (Miltenyi Biotec). Suspensions were filtered through 40 μM strainer and rinsed with MACS buffer. The pellet was resuspended in RBC lysis buffer, washed with MACS, and incubated with CD45 magnetic beads to deplete CD45+ cells. CD45-cell fraction was collected using the MultiMACS Cell24 Separator (Miltenyi Biotec). Cells were sorted as described above. Following sorting, 5×10^3^ zsGreen+ AT2 were mixed with 5×10^4^ fibroblasts in 50% Matrigel (Corning, 356231) and 50% organoid media (αMEM, penicillin/streptomycin, insulin/transferrin/selenium, FBS, heparin, Amphotericin B, Rock1 inhibitor, L-glutamine) in 0.4 μM Transwell inserts. Cell/Matrigel mixtures were solidified at 37°C for 5 minutes before 300μl of media was added under the Transwell. 24 hours after isolation, organoids received media change every other day containing either 1 μM ISRIB or DMSO for 10 days.

#### Tissue sectioning, immunofluorescence and image analysis

After euthanasia and perfusion, trachea was cannulated with Luer syringe stub blunt needle and mouse lungs were inflated with 4% paraformaldehyde at 15 cm H_2_O column pressure. Lungs were fixed in 4% paraformaldehyde (PFA) overnight at 4°C followed by cryo-embedding in optimal cutting-temperature compound (OCT). Sections were cut at 14μM on a cryostat. For immunostaining, sections were blocked in 10% normal goat serum (NGS), 0.3%TritonX-100 in PBS for 30 minutes followed by an additional 1:10 anti-mouse Fab (H+L) for 2 hours. Primary antibodies were prepared in 2% NGS 0.3%TritonX-100 PBS and applied to sections overnight at 4°C. Sections were washed in PBS followed by secondary antibodies for 30 minutes. Lastly, sections were incubated in Hoechst (1:10,000) for 5 minutes followed by a brief incubation of 0.3% Eriochrome black prior to mounting. Primary antibodies were used at concentrations described below. *For 2D culture*, cells were fixed 4 days after isolation in 4% PFA. For the alveolar organoids, organoids were fixed 10 days after isolation. Fixed cells and organoids were neutralized in 10 mM glycine followed by blocking in 10% normal goat serum (NGS) in 0.3%TritonX-100 PBS. Primary antibodies were prepared in 3% NGS in 0.3%TritonX-100 PBS and applied for 1 hour at room temperature. Following PBS washes, fluorophore-conjugated secondary antibodies were prepared at 1:300 for 30 minutes in the dark at room temperature. Hoechst was diluted 1:10,000 and applied for 5 minutes at room temperature following secondary antibody incubation. Coverslips and filters were mounted using ProLong Gold anti-fade solution. Images were obtained using Zeiss Axioplan epifluorescence and Nikon W1 Spinning Disk confocal microscopes. *For nuclear area quantification*, 5 fields of view were selected for each condition per biological replicate. Images were taken on a Zeiss microscope and analyzed in Fiji using the Stardist plugin. Primary antibodies were as follows: SFTPC (1:100); beta-catenin (1:100); Podoplanin (1:30); Krt8 (1:30), and F-actin (1:50). *For thick section imaging with tissue clearing*, lungs were sliced in PBS using a vibratome (Leica VT1200s) in 200μm thick sections. Tissue slices were cleared using the LifeCanvas Technologies SHIELD method (Cambridge, MA, USA). *For 3-D volumetric analysis*, 200 μM thick sections were imaged sequentially in 0.5 μM steps on a Nikon W1 Spinning Disk confocal microscope. Cell volumes were measured using AT2-lineage label (YFP) via Imaris.

#### Live-cell imaging

Freshly isolated alveolar epithelial cells were plated at 500K/well in a 12 well plate. 48 hours post-isolation, media was changed and adherent cells were imaged every 15-minutes on a Biostation-CT system (Nikon Instruments, New York, USA) for a total of 5 hours. Image sequences were analyzed in Fiji (ImageJ v: 2.1.0/1.53c).

#### Statistical analysis

Data analysis and statistical tests were performed using GraphPad Prism software (v.9.2.0).

## Acknowledgements

This work relied on the following Northwestern University services and core facilities: Flow Cytometry (NCI CA060553, 1S10OD011996, 1S10OD026814); Center for Advanced Microscopy (NCI CCSG P30 CA060553, NCRR 1S10 RR031680, 1S10OD021704); BioCryo facility of Northwestern University’s NUANCE Center (NSF ECCS-2025633, DMR-1720139).

## Author contributions

AW, MMH, SW, ASF, LW and RPA designed and conducted experiments; AW, MMH, ASF, AM and CJG analyzed results; RG wrote a macro for image analysis; AM advised and analyzed flow cytometry, SH, LD, MH, AM, GRSB advised and discussed results; CJG designed and supervised study, performed analysis and wrote manuscript. MH, AM, GRSB and CJG provided funding for project.

---

**Figure S1:**
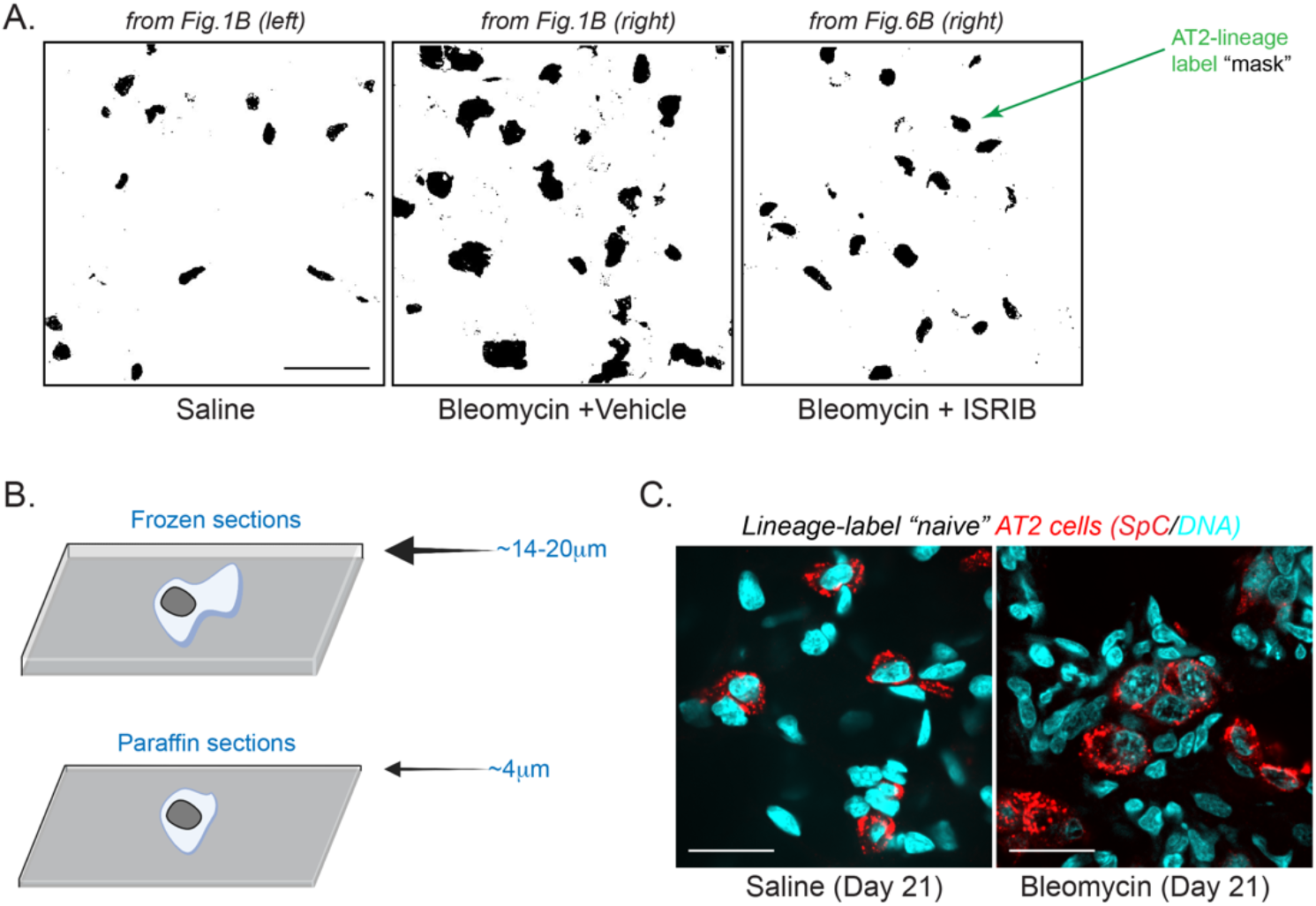
AT2 cell hypertrophy by frozen- and thick-section image analysis. (**A**) Black and white mask of images in Figure 1 for quantification of lineage-labelled AT2 cell area and perimeter (black objects, green arrow). (**B**) Injury-induced AT2 hypertrophy readily observed by frozen sections, which are ~4-fold thicker than typical paraffin sections. (**C**) Hypertrophic AT2 phenotype is not an artefact of lineage-labeling AT2 cells. Confocal image of Vibratome-cut 200μm thick lung section from a C57BL/6 mouse with clearing (SHIELD method) immunofluorescence detection of SpC (red) and nuclei (Draq5; cyan). Note bleomycin-treated AT2 cells are clearly larger, even 21 days post-bleomycin instillation.

**Figure S2:**
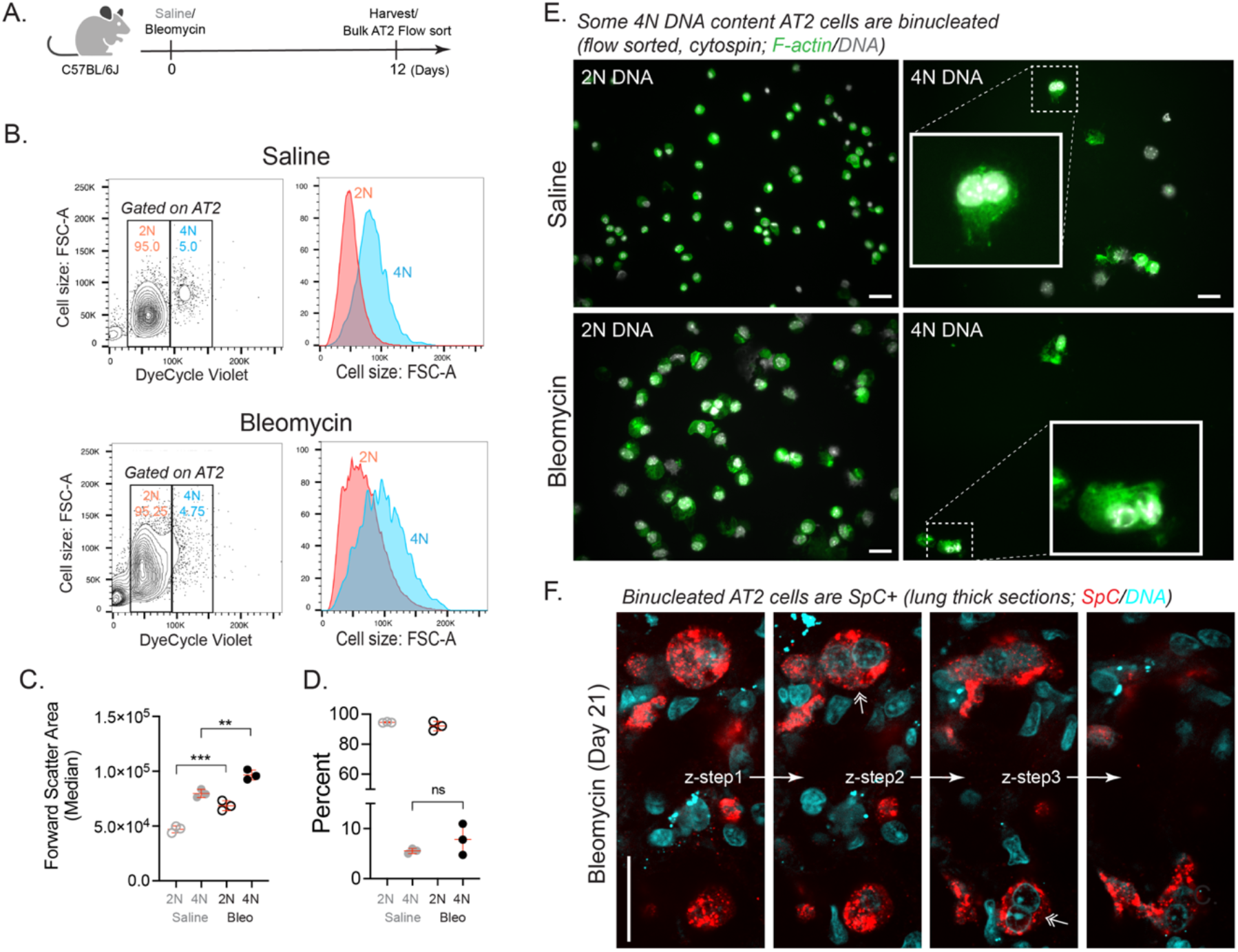
Quantification of injury-induced AT2 hypertrophy and binucleation by flow cytometry. (**A**) Schematic of experimental design. (**B**) Young adult mice (3-5 months) were administered saline alone or 0.025 units of bleomycin and lungs were harvested and digested for flow cytometry on Day 12 with representative flow plots gated on CD45-/EpCAM+ AT2 cells with 2N (diploid, red) or 4N DNA content (cyan) (Methods). (**C**) Quantification of AT2 cell size (median Forward Scatter Area). (**D**) Percent 2N versus 4N AT2 cells. N=3 mice. p=0.0008 and 0.0039 by ANOVA with Tukey’s. (**E**) Validation of bleomycin-induced AT2 cell size increase and binucleation by cytospin analysis. Cellular F-actin stained with phalloidin (green) to distinguish *bona fide* binucleation, with absence of F-actin-rich cell-cell junction between nuclei. (**F**) Binucleated AT2 cells can be SpC+. Confocal z-stack image of Vibratome-cut 200μm thick lung section from a C57BL/6 mouse with clearing (SHIELD method) immunofluorescence detection of SpC (red) and nuclei (Draq5; cyan). Double arrowhead implicates two binucleated AT2 cells in the same field of view.

**Figure S3:**
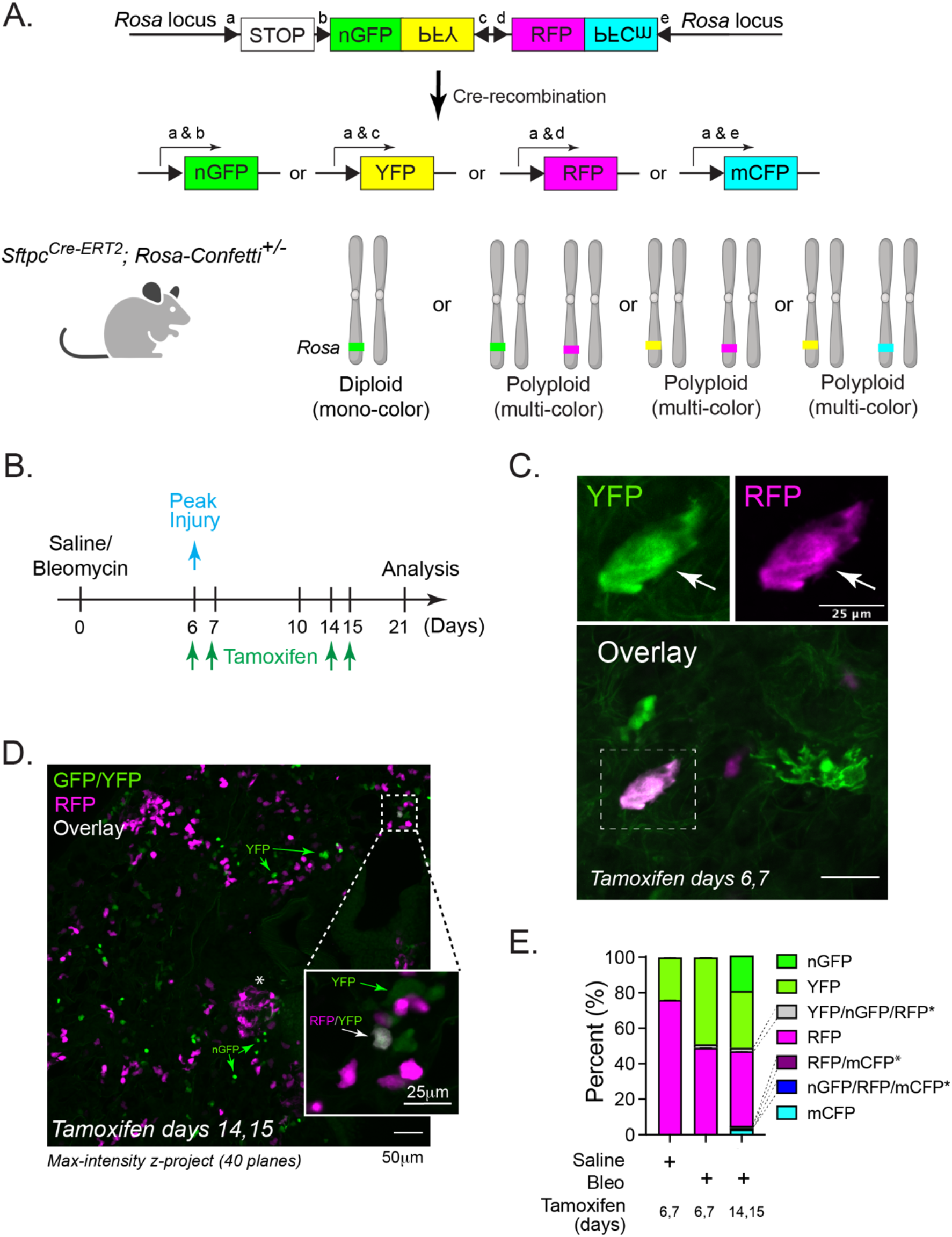
Identification of rare ≥4N AT2 cells via stochastic multi-color Confetti-labeling system. (**A**) Schematic adapted from (39) using *Sftpc*^Cre-ER; R26R-Confetti^ strain (14). (**B**) Tamoxifen activation of *Sftpc*-targeted lineage-label at days 6-7 post-bleomycin injury and harvested Day 21. (**C**) Confocal image of thick-section cleared (SHIELD) lung, maximum z-projection of 3 slices showing rare YFP (green)/RFP (magenta) dual-positive cells, suggesting mononuclear polyploidy from partial nuclear exclusion of signal. The enlarged/flattened size of this cell (>25μm) is also compatible with 4N polyploidy. (**D**) Lower magnification view maximum z-projection (40 planes) shows that most lineage labeled cells are either nuclear GFP (nGFP), YFP (cytoplasmic) or RFP positive, with very few nGFP or YFP and RFP-dual-positive cells (see inset showing gray cell, which reflects overlay of YFP (green) and RFP (magenta) signals). As this cell is ~10μm, it likely reflects a G2-phase AT2 cell with 4N DNA content. Asterisk validates bleomycin injury showing RFP lineage label along flattening AT1 cells. (**E**) Quantification of C-D from n=2 mice/condition. Note our microscopes do not discriminate between nGFP/YFP signals, due to spectral overlap of these fluorescent proteins. Fortunately, distinguishing between nGFP/YFP is not critical to answer the polyploidy question—as nGFP or YFP *<u>and</u>* RFP is sufficient evidence of >2N DNA content, where we see this combination only rarely (~1-2%). Few nGFP cells are detected when we activate the lineage label shortly after peak injury (Days 6, 7 versus 14, 15), possibly due to nuclear nGFP toxicity (74). Lastly, CFP cells are typically detected less frequently with this reporter, possibly due to its placement in the cassette (39).

**Figure S4:**
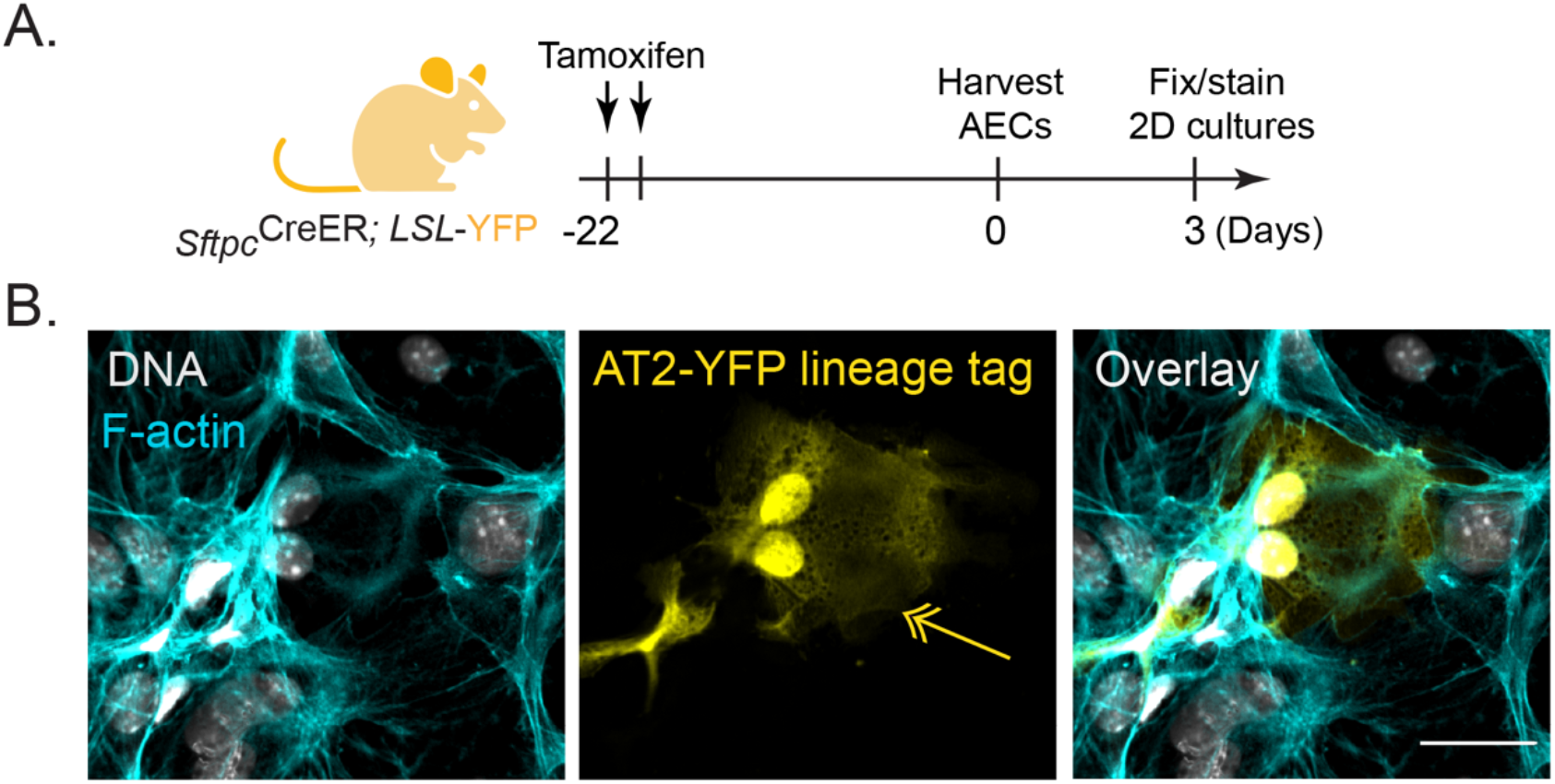
Detection of binucleated, lineage-labeled mouse AT2 cells in cell culture. (**A**) Schematic for activation of AT2-lineage label before AT2 isolation. (**B**) Confocal image of lineage-labeled AT2 cell (yellow) grown 3 days on glass. Arrow indicates binucleated, AT2-derived cell. F-actin in cyan.

**Figure S5:**
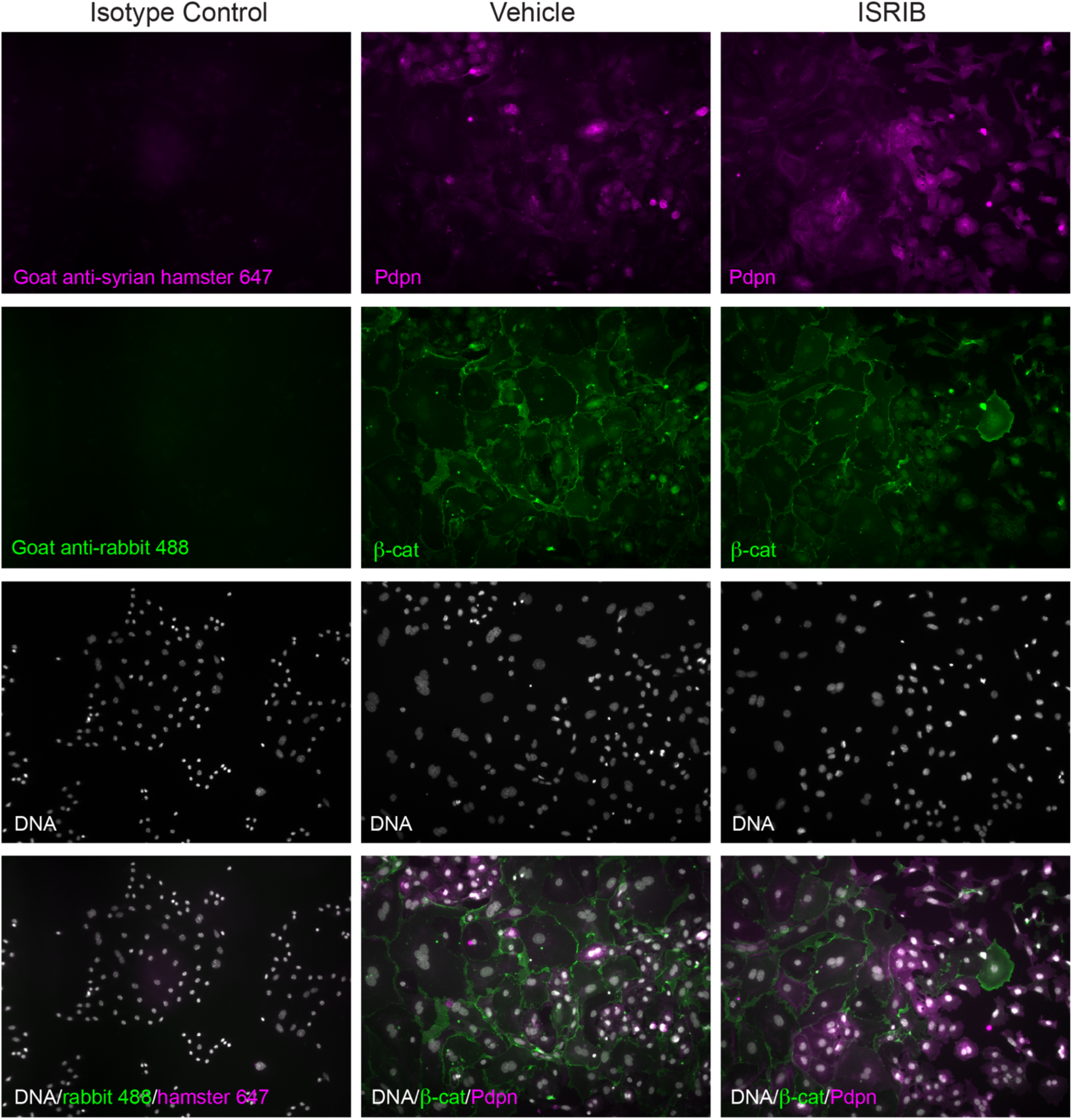
Antibody-isotype control to validate podoplanin staining in ex vivo mouse AT2 cell cultures. Individual fluorescence panels from overlay images in Figure 6D.

